# Hearing in categories aids speech streaming at the “cocktail party”

**DOI:** 10.1101/2024.04.03.587795

**Authors:** Gavin M. Bidelman, Fallon Bernard, Kimberly Skubic

**Author notes:** **Author contributions:** G.M.B. designed the experiment, F.B. and K. S. collected the data, G. M. B., F.B., and K. S. analyzed the data, and all authors wrote the paper. **Address for editorial correspondence:** Gavin M. Bidelman, Ph.D., Speech, Language and Hearing Sciences 2631 East Discovery Parkway Bloomington, IN 47408.

## Abstract

Our perceptual system bins elements of the speech signal into categories to make speech perception manageable. Here, we aimed to test whether hearing speech in categories (as opposed to a continuous/gradient fashion) affords yet another benefit to speech recognition: parsing noisy speech at the “cocktail party.” We measured speech recognition in a simulated 3D cocktail party environment. We manipulated task difficulty by varying the number of additional maskers presented at other spatial locations in the horizontal soundfield (1-4 talkers) and via forward vs. time-reversed maskers, promoting more and less informational masking (IM), respectively. In separate tasks, we measured isolated phoneme categorization using two-alternative forced choice (2AFC) and visual analog scaling (VAS) tasks designed to promote more/less categorical hearing and thus test putative links between categorization and real-world speech-in-noise skills. We first show that listeners can only monitor up to ∼3 talkers despite up to 5 in the soundscape and streaming is not related to extended high-frequency hearing thresholds (though QuickSIN scores are). We then confirm speech streaming accuracy and speed decline with additional competing talkers and amidst forward compared to reverse maskers with added IM. Dividing listeners into “discrete” vs. “continuous” categorizers based on their VAS labeling (i.e., whether responses were binary or continuous judgments), we then show the degree of IM experienced at the cocktail party is predicted by their degree of categoricity in phoneme labeling; more discrete listeners are less susceptible to IM than their gradient responding peers. Our results establish a link between speech categorization skills and cocktail party processing, with a categorical (rather than gradient) listening strategy benefiting degraded speech perception. These findings imply figure-ground deficits common in many disorders might arise through a surprisingly simple mechanism: a failure to properly bin sounds into categories.

## 1. Introduction

Perceptual organization requires sensory phenomena be subject to invariance: features are mapped to common equivalencies by assigning similar objects to the same category membership (Goldstone and Hendrickson, 2010). Categories occur in all aspects of human cognition including face (Beale and Keil, 1995), color (Franklin et al., 2008), and music (Locke and Kellar, 1973; Siegel and Siegel, 1977; Burns and Ward, 1978; Zatorre and Halpern, 1979; Howard et al., 1992; Klein and Zatorre, 2011) perception. But categories are particularly important in the context of spoken word recognition (Liberman et al., 1967; Pisoni, 1973; Harnad, 1987; Pisoni and Luce, 1987). In speech perception, categories help bootstrap comprehension by generating perceptual constancy in the face of acoustic variability (e.g., talker variation, signal corruption) (Prather et al., 2009). Thus, hearing in categories might help bolster speech-in-noise (SIN) skills by constraining and reducing perceptual variability in the speech signal.

Indeed, emerging evidence suggests that forming categories might benefit speech perception in noisy environments. In naturalistic soundscapes, the auditory system must extract target speech and simultaneously filter out extraneous sounds in what is described as the “cocktail-party problem” (Cherry, 1953; Bregman, 1978; Yost, 1997). Theoretically, once equivalency between stimuli is formed, irrelevant variations among them can be deemphasized (Goldstone and Hendrickson, 2010). Based on this premise, we have hypothesized that hearing speech in a categorical mode (a more abstract level of coding) might help aid degraded speech perception since irrelevant variations in the physical surface features of the signal can be largely discarded in favor of retaining a more abstract, phonetic code for speech (see Bidelman et al., 2020). Supporting this notion, we have recently shown speech categories are surprisingly robust to acoustic interference, diminishing only at severe noise levels [i.e., negative signal-to-noise ratios (SNRs)] (Bidelman et al., 2019; Bidelman et al., 2020; Lewis and Bidelman, 2020; Bidelman and Carter, 2023). These behavioral results are bolstered by neuroimaging data which reveal the brain’s encoding of speech is not only enhanced for sounds carrying a clear phonetic identity compared to their phonetically ambiguous counterparts but that category members are actually more resistant to external acoustic noise (Gifford et al., 2014; Bidelman et al., 2020). Similar parallels are found in the visual domain (Helie, 2017).

Further support for the link between categorical/discrete hearing modes of listening and SIN processing stems from studies in both highly skilled and impoverished listeners. For example, musicians demonstrate improved figure-ground perception in a variety of SIN tasks (Bidelman and Krishnan, 2010; Parbery-Clark et al., 2011; Swaminathan et al., 2015; Anaya et al., 2016; Clayton et al., 2016; Brown et al., 2017; Deroche et al., 2017; Du and Zatorre, 2017; Mankel and Bidelman, 2018; Torppa et al., 2018; Yoo and Bidelman, 2019), as well as better multi-talker streaming (Bidelman and Yoo, 2020). Musicians also show enhanced categorization for speech and musical sounds in the form of more discrete, binary labeling of tokens along graded continua (Bidelman et al., 2014; Bidelman and Alain, 2015; Bidelman and Walker, 2019). On the contrary, several clinical populations involving auditory-based and learning disorders (e.g., dyslexia) can show weaker phoneme categorization (Thibodeau and Sussman, 1979; Jerger et al., 1987; Werker and Tees, 1987; Noordenbos and Serniclaes, 2015; Gabay et al., 2023) and poorer SIN processing (Cunningham et al., 2001; Warrier et al., 2004; Putter-Katz et al., 2008; Lagacé et al., 2010; Dole et al., 2012; Dole et al., 2014) than their normally developing peers. The neural basis of acoustic-phonetic processing depends on a strong auditory-sensory memory interface (DeWitt and Rauschecker, 2012; Bizley and Cohen, 2013; Chevillet et al., 2013; Jiang et al., 2018) rather than higher-level cognitive faculties (e.g., attentional switching and IQ; Kong and Edwards, 2016). Thus, the degree to which listeners show categorical (discrete) vs. gradient (non-categorical) perception could have ramifications for understanding clinical disorders that impair SIN processing. A failure to flexibly warp acoustic representations of the speech signal into well-formed, discrete categories could provide a linking hypothesis to describe individual differences in perceptual SIN skills among normal and clinical populations alike.

Conversely, an alternate view argues that gradient/continuous listening strategies might help facilitate SIN processing. Under this notion, maintaining sensitivity to within-category information (and even nuisance details of the noise itself) might allow more nimble perceptual readout of speech information (Kapnoula et al., 2017; Apfelbaum et al., 2022). In other words, higher sensitivity to within-category information could offer more flexible processing, allowing listeners to “hedge” their bets in the face of ambiguity (Kapnoula et al., 2017). However, when tested empirically, gradient (non-categorical) perception is generally *not* associated with speech-in-noise listening performance (Kapnoula et al., 2017; Kapnoula et al., 2021), suggesting that while listeners do have simultaneous access to continuous, within-category cues (Pisoni and Lazarus, 1974; Pisoni and Tash, 1974; Spivey et al., 2005; Huette and McMurray, 2010; Bidelman and Carter, 2023), they do not readily exploit them when parsing speech in ambiguous or degraded conditions (cf. Kapnoula et al., 2017). Instead, both the construction of discrete perceptual objects and natural binning process of categorization might better enable category members to “pop out” among a noisy feature space, thereby facilitating SIN processing (e.g., Nothdurft, 1991; Pérez-Gay Juárez et al., 2019; Bidelman et al., 2020). This premise lays the groundwork for testing critical but yet undocumented links between categorization and SIN listening skills. These concepts have enjoyed long but separate histories in the literature which we now bridge in the current study.

To this end, we measured speech-in-noise processing and phonetic categorization in young, normal hearing listeners to assess putative relations between these fundamental skills in speech perception. Because SIN perception also relates to high-frequency hearing sensitivity even in “normal hearing” individuals (Monson et al., 2019; Mishra et al., 2022), we also measured extended high-frequency (EHF) audiometric thresholds. Noise-degraded speech perception abilities were assessed using standard clinical (i.e., QuickSIN; Killion and Niquette, 2000) and ecological SIN assays. For the latter, we used a simulated, multi-talker cocktail party environment to assess real-world SIN perception abilities that engage auditory segregation and streaming processes (Bidelman and Yoo, 2020). Performance on this task is largely independent of cognitive factors including sustained attention, working memory, and IQ, suggesting it has high construct validity and is not easily explainable by mere cognitive differences between listeners (Bidelman and Yoo, 2020). Participants monitored target sentences [Coordinate Response Measure (CRM) corpus] (Bolia et al., 2000) presented simultaneous with up to 4 additional talkers (other CRM sentences). Critically, we presented masking talkers in either a forward or time-reversed direction to induce more/less informational masking (IM). Forward masks were predicted to be more difficult since they are clearly recognized as speech carrying linguistic information and this should interfere with target recognition (i.e., increased informational, speech-on-speech masking). The time-reversal in reversed masks, on the other hand, destroys their lexical information and was expected to produce only energetic masking on the target—making the task easier. The difference between conditions was used to index cocktail party performance, i.e., the degree to which a listener experiences the added interference of IM (Swaminathan et al., 2015; Yoo and Bidelman, 2019).

Categorization for labeling isolated acoustic-phonetic speech sounds was measured using two different continua (vowels vs. CVs) presented under different task structures (two alternative forced choice—2AFC vs. visual analog scale—VAS). These manipulations allowed us to assess categorization under stimulus and task conditions designed to promote discrete (2AFC) vs. gradient (VAS) hearing, respectively. CVs are perceived more categorically than vowels (Pisoni, 1973; Altmann et al., 2014; Carter et al., 2022) and binary responding (2AFC) produces stronger categorical hearing during labeling than classifying the same speech sounds using a VAS scale (Kapnoula et al., 2017). Relevant to the current study, VAS categorization has been used to measure the degree of categoricity in a listener’s perception, since it allows for more graded judgments of the acoustic-phonetic space than a binary 2AFC task. Importantly, the VAS approach can identity listeners that respond in a discrete (categorical) vs. gradient (continuous) manner (Kapnoula et al., 2017). Based on prior work (Bidelman et al., 2019; Bidelman et al., 2020; Lewis and Bidelman, 2020; Bidelman and Carter, 2023), we hypothesized that more categorical listeners (i.e., more binary responders) would show more successful QuickSIN and/or cocktail party streaming perception. Alternatively, if a continuous listening strategy is more beneficial for SIN processing (Kapnoula et al., 2017), more graded responders in VAS phoneme labeling should show improved SIN performance. To anticipate, our findings establish a categorization-SIN link whereby more discrete (rather than gradient) categorization benefits degraded cocktail party speech perception.

## 2. Materials & Methods

### 2.1 Participants

N=21 young (age range: 22-37 years; 9 male, 12 female), normal-hearing adult participants were recruited for the study. On average, they had 18 ± 1.1 years of education and were right-handed (72.6 ± 39.9 % handedness laterality; Oldfield, 1971). All showed normal hearing sensitivity (puretone audiometric thresholds ≤25 dB HL, 250 to 20000 Hz; see **Fig. 2**). Non-native speakers perform worse on SIN tasks than their native-speaking peers (Rogers et al., 2006; Bidelman and Dexter, 2015). Thus, all participants were required to be native English speakers. The sample was largely “nonmusicians,” averaging 6.6 ± 6.2 years of formal music training (Wong et al., 2007; Parbery-Clark et al., 2009; Yoo and Bidelman, 2019; MacLean et al., 2024). It should be noted that >10 years of music engagement is generally needed before observing musician-related benefits in SIN (Parbery-Clark et al., 2009; Yoo and Bidelman, 2019) or cocktail party speech perception (Bidelman and Yoo, 2020). Indeed, participants’ years of musical training was not correlated with any of the dependent variables (all *p*s > 0.05). Each participant gave written informed consent in accordance with a protocol approved by the University of Memphis Institutional Review Board.

**Figure 1:**
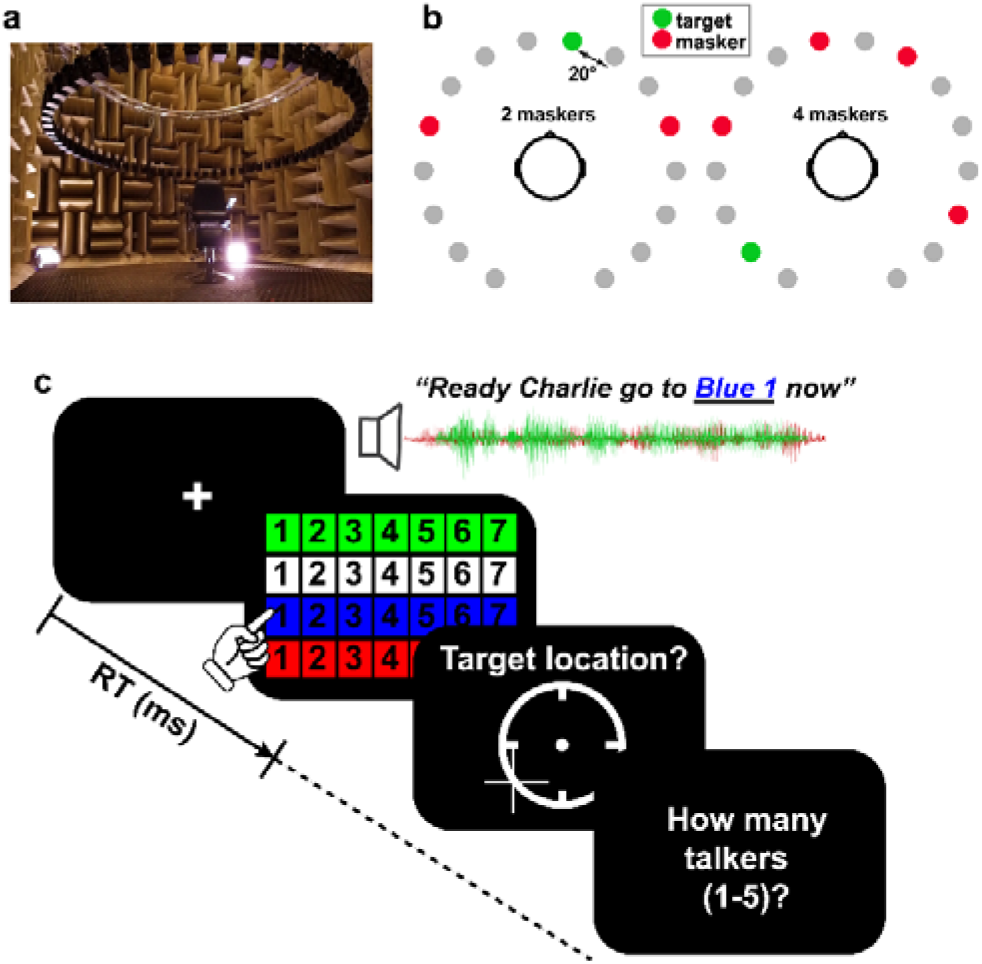
Cocktail party streaming task. (**a**) Participants were seated in the center of a 16-ch speaker array within an anechoic chamber. Speaker heights were positioned at ear level (∼130 cm) during the task with a radial distance of 160 cm to the center of the head and speaker-to-speaker distance of ∼20^0^. **(b**) Example stimulus presentation (2 and 4 masker talker conditions). Participants were asked to recall the color, number, and perceived location of target callsign sentences from the CRM corpus (Bolia et al., 2000). Target location was varied randomly from trial to trial and occurred simultaneously with between 0 and 4 concurrent talkers presented in either forward or time-reversed directions. (**c**) Example trial time course. After presentation of CRM sentences, listeners recalled the color-number combination of the target talker, its perceived location in the hemifield, and how many talkers they heard in the soundscape.

**Figure 2:**
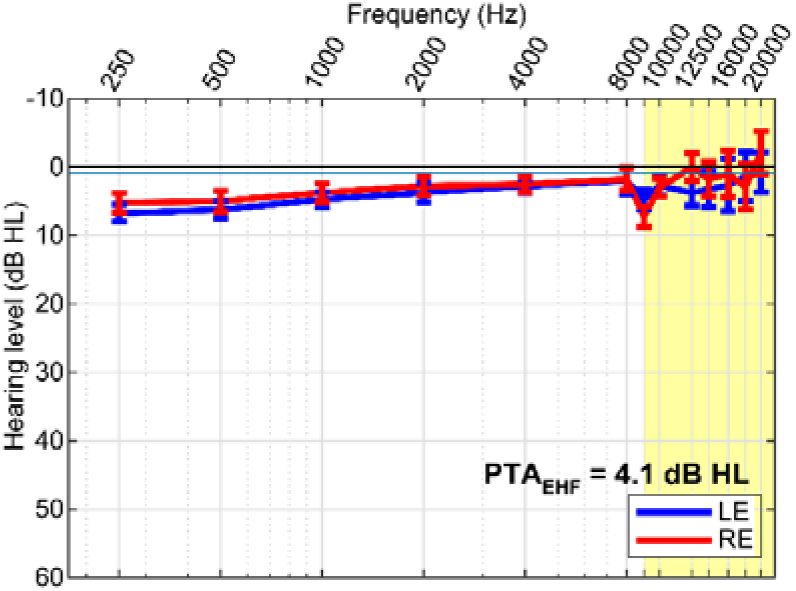
Extended high frequency (EHF) hearing thresholds. Audiograms for left (LE) and right (RE) ears. Pure-tone average (PTA) EHF thresholds in the normal and EHF (9-20 kHz; yellow highlight) frequency range were well within normal hearing limits. errorbars = ± 1 s.e.m.

### 2.2 Stimuli & task paradigms

#### Speech streaming task

We measured naturalistic cocktail party listening skills via a sentence-on-sentence speech recognition task conducted in a 3D spatial soundfield (Bidelman and Yoo, 2020). Speech recognition and localization performance was assessed in a simulated multi-talker cocktail party environment within the University of Memphis Anechoic Chamber (**Fig. 1a**)^1^. A 16-channel circular speaker array was positioned vertically 130 cm above the mesh floor of the anechoic chamber (approximately ear height). Each speaker had a radial distance of 160 cm to the center of the head. Speaker-to-speaker distance was ∼20 degrees. Stimuli were presented at 70 dB SPL (z-weighted, freefield), calibrated using a Larson–Davis sound level meter (Model LxT).

We used Coordinate Response Measure (CRM) sentences (Bolia et al., 2000) to measure speech recognition in a multi-talker sounds mixture. CRM sentences contain a different target callsign (Charlie, Ringo, Laker, Hopper, Arrow, Tiger, Eagle, Baron), color (Blue, Red, White Green), and number (1-8) combination embedded in a carrier phrase (e.g., “Ready <u>Charlie</u>, go to <u>blue three</u> now”). The corpus contained all possible permutations of these callsign-color-number combinations spoken by eight different talkers (male and female). We used CRM sentences as they are sufficiently novel to listeners to avoid familiarity effects that might confound SIN recognition (Johnsrude et al., 2013; Holmes et al., 2018; Brown and Bidelman, 2022). They are also natural productions that offer a level of control (e.g., similar length, same sentence structure). Participants were cued to the target callsign before each block and were instructed to recall its color-number combination via a computer screen GUI as fast and accurately as possible (e.g., “b2” = blue-two; “r6” = red-six; **Fig. 1c**). We logged both recognition accuracy and reaction times (RTs). RTs were clocked from the end of the stimulus presentation.

On each trial, listeners heard a mixture of sentences one of which contained the target callsign and additional CRM sentence(s) that functioned as multi-talker masker(s). Three additional constraints were imposed on sentence selection to avoid unnecessary task confusion: (1) targets were always from the same talker and callsign (within a block); (2) maskers were absent of any callsign, color, and number used in the target phrase (i.e., the callsign’s information was unique among the speech mixture); (3) target and masker(s) were presented from unique spatial locations (i.e., different speakers). The target speaker/callsign was allowed to vary between blocks but was fixed within block. Male and female talkers were selected randomly. Thus, on average, targets and maskers were 50% male and 50% female. Presentation order and spatial location of the sentences in the 360-degree soundfield were otherwise selected randomly (**Fig. 1b**).

We manipulated task difficulty by parametrically varying the number of additional maskers on a trial-by-trial basis (0, 1, 2, 3, 4) presented at other spatial locations in the speaker array. All talkers were presented with an equivalent level (i.e., RMS amplitude). We required participants to identify *both* the call color and number of the target callsign phrase to be considered a correct response (chance level = 3.13% = 1/32). It is possible for listeners to localize sound sources even if they cannot identify them (Rakerd et al., 1999). Consequently, after recognition, we had participants indicate the perceived location (azimuth) of the target by clicking on a visual analogue of the speaker array displayed on the screen. Lastly, listeners indicated the number of total talkers they perceived in the soundfield to gauge source monitoring abilities (Yost et al., 2019). An example trial timecourse is shown in **Fig. 1c**.

This identical CRM task was run in two masking conditions: (i) forward and (ii) time-reversed maskers (random order). Forward maskers consisted of the CRM sentences unmanipulated. In the reverse condition, the masking talkers were time-reversed. These two conditions allowed us to assess streaming during informational (forward) vs. energetic (reverse) acoustic interference while controlling for the SNR and spectrotemopral characteristics of the maskers (Carter and Bidelman, 2021). There were total of 32 trials per noise block, repeated twice (i.e., 64 trials per masker condition).

#### Phoneme categorization

##### Vowel and CV continua

The vowel continuum was a synthetic 5-step vowel continuum spanning from “u” to “a” (Bidelman et al., 2014; Bidelman and Walker, 2017; Bidelman et al., 2020; Carter et al., 2022). Each token was separated by equidistant steps acoustically based on first formant frequency (F1). Individual tokens were 100 ms in duration including 5 ms of ramping. Each contained identical voice fundamental (F0), second (F2), and third formant (F3) frequencies (F0: 150, F2: 1090, and F3: 2350 Hz), chosen to roughly approximate productions from male speakers (Peterson and Barney, 1952). F1 was parameterized over five equal steps between 430 and 730 Hz such that the resultant stimulus set spanned a perceptual phonetic continuum from /u/ to /a/ (Bidelman et al., 2013).

The consonant vowel (CV) continuum consisted of a 5-step, stop-consonant /da/ to /ga/ sound gradient (varying in place of articulation) (e.g., Bidelman et al., 2019; Carter et al., 2022). Original speech utterances were adopted from Nath and Beauchamp (2012). Individual tokens were 350 ms in duration including 5 ms of ramping. Stimulus morphing was achieved by altering the F2 formant region in a stepwise fashion using the STRAIGHT software package (Kawahara et al., 2008).

##### 2AFC vs. VAS categorization task

Categorization for both continua were measured under two task paradigms: (i) 2 alternative-forced choice (2AFC) binary key press or (ii) mouse click on a visual analog scale (VAS) (Massaro and Cohen, 1983; Kong and Edwards, 2016; Kapnoula et al., 2017) (see **Fig. 4, *insets***). 2AFC and VAS tasks were run in separate (randomized) blocks but used otherwise identical speech stimuli; only the task paradigm differed. The VAS paradigm required participants to click a point along a continuous visual scale with endpoints labeled “u”/”da” and “a”/”ga” to report their percept. Use of the entire analog scale was encouraged. Unless the participants had clarifying questions, no other instructions were provided (Kapnoula et al., 2017).

**Figure 3:**
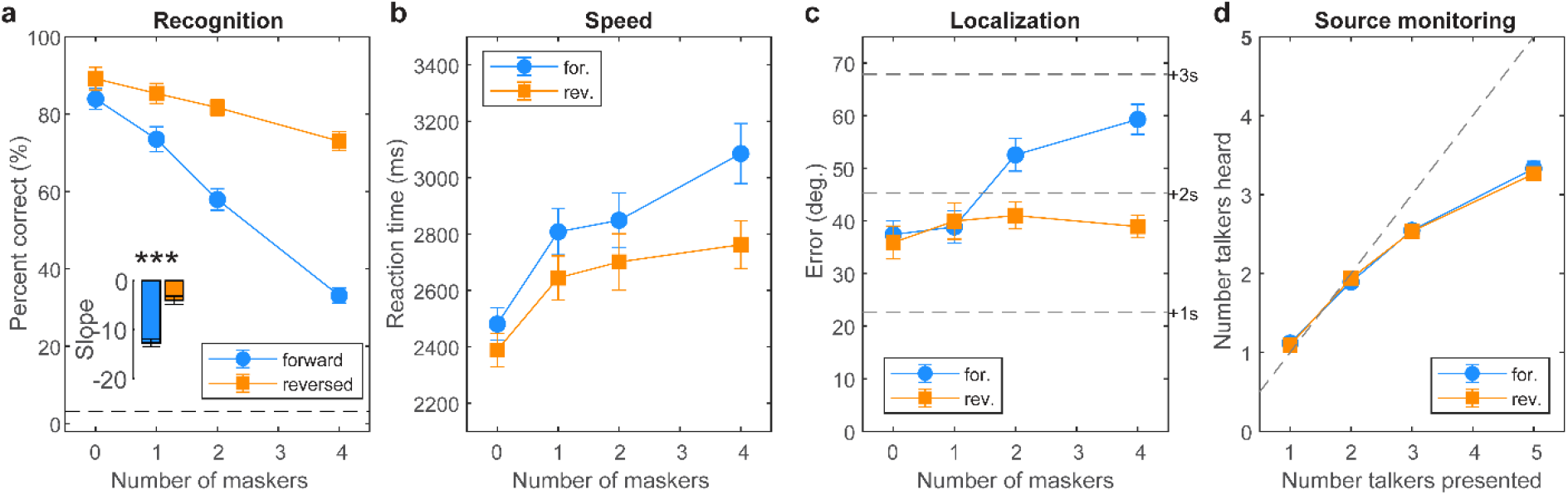
Cocktail party listening performance. (**a**) Speech recognition declines with increasing masker counts but is much poorer under informational/linguistic vs. purely energic masking (cf., forward vs. reverse masker directions) (inset). Dotted line = chance performance. (**b**) Forward maskers yield slower recognition speeds than the reverse maskers owing to their added linguistic interference. (**c**) Listeners localized targets within 2 speakers (40-60° error) with better localization during purely energetic masking. (**d**) Source monitoring. Listeners saturate in source monitoring and only report hearing up to ∼3 additional talkers despite up to 5 in the soundscape. errorbars = ± 1 s.e.m., \*\*\**p*<0.0001

**Figure 4:**
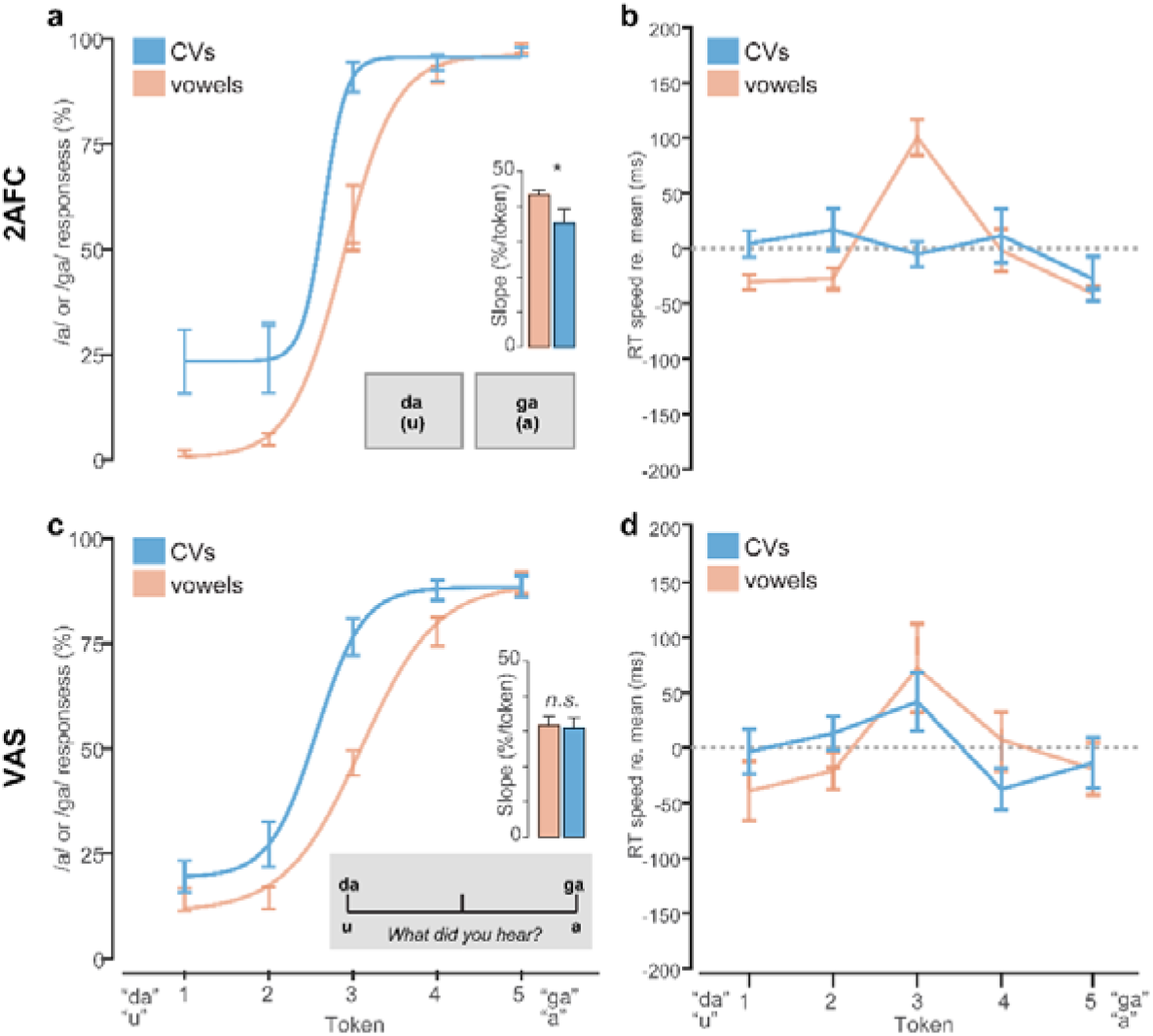
Stimulus- and task-dependent changes in the strength of perceptual categorization. Speech categorization and RT speeds under (**a-b**) 2AFC and (**c-d**) VAS labeling tasks. Note the sharper, more discrete categorization for CVs compared to vowels in the 2AFC (but not VAS) condition. RTs show the typical slowing near the perceptually ambiguous midpoint of the vowel (but not CV) continuum for both tasks. RTs are normalized to the global mean to highlight token-and stimulus-related changes. However, VAS responses were 750 ms slower than 2AFC across the board. errorbars = ± 1 s.e.m., \**p*<0.05.

Speech stimuli were delivered binaurally at a comfortable listening level (∼75 dB SPL) through Sennheiser HD 280 circumaural headphones. Listeners heard 75 trials of each individual speech token (per task block). On each trial, they were asked to label the sound with a response (“u” or “a”; “da” or “ga”) as quickly and accurately as possible. Following listeners’ behavioral response, the interstimulus interval (ISI) was jittered randomly between 800 and 1000 ms (20 ms steps, uniform distribution) to avoid anticipation of subsequent stimuli. In total, there were four categorization conditions: /u/-/a/ and /da/-/ga/ continua presented in either a 2AFC or VAS paradigm.

#### QuickSIN

The QuickSIN (Killion et al., 2004) provided a normed test of SIN perceptual abilities. Participants heard six sentences embedded in four-talker noise babble, each containing five keywords. Sentences were presented at 70 dB HL. The signal-to-noise ratio (SNR) decreased parametrically in 5 dB steps from 25 dB SNR to 0 dB SNR. At each SNR, participants were instructed to repeat the sentence and correctly recalled keywords were logged. We computed their SNR loss by subtracting the number of recalled target words from 25.5 (i.e., SNR loss = 25.5-Total Correct). The QuickSIN was presented binaurally via Sennheiser HD 280 circumaural headphones. Two lists were run and the second was used in subsequent analysis to avoid familiarization effects (Yoo and Bidelman, 2019; Bidelman and Yoo, 2020).

#### Extended high-frequency (EHF) thresholds

In addition to standard pure-tone air-conduction audiometry, we measured hearing thresholds at EHFs of 9, 10, 12.5, 14, 16, 18, 20 kHz. EHFs were measured using circumaural headphones (Sennheiser HDA 200, Wedemark, Germany) specialized for high-frequency audiometry.

### 2.3 Statistical analysis

Unless otherwise noted, we analyzed the dependent variables using mixed-model ANOVAs in R (version 4.2.2) (R-Core-Team, 2020) and the lme4 package (Bates et al., 2015). Speech streaming measures (%-accuracy, RTs, localization error, source monitoring) were analyzed with fixed effects of masker count (0-4) and masker direction (forward, reverse). Phoneme categorization measures (identification slope, RTs) were analyzed with fixed effects of task (2AFC, VAS), continuum (vowels, CVs), and—in the case of RTs—token (Tk1-5). Subjects served as a random effect. Tukey-adjusted contrasts were used for multiple comparisons. %-correct data were RAU transformed prior to statistical treatment (Studebaker, 1985). Effect sizes are reported as partial eta squared (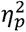) and degrees of freedom (*d.f.*) using Satterthwaite’s method.

## 3. Results

### 3.1 High-frequency thresholds

Grand average extended high-frequency (EHF) audiometric thresholds are shown for the left and right ear in **Figure 2**. EHFs in the 9-20 kHz frequency range were unremarkable and within normal limits for all listeners (average PTA_9-20kHz_ = 4.1 ± 10.5 dB HL).

### 3.2 “Cocktail party” speech streaming

Streaming performance measures (i.e., %-accuracy, RTs, localization error, source monitoring) are shown in **Figure 3**. Speech recognition expectedly declined from ceiling to near-floor performance with increasing masker counts from 0 (unmasked) to 4 multi-talkers. Still, all listeners showed above-chance recognition even amidst 4 maskers (all *p*s< 0.0001; *t*-test against 3.13% chance). Notably, we found a masker direction x masker count interaction on target speech recognition accuracy [*F_3,140_* =7.93, *p*<0.0001, 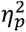*=* 0.15; **Fig. 3a**]. The interaction was attributable to a stronger decline in speech recognition performance with increasing talkers amidst forward compared to reversed maskers (Fig. 3a, *inset*; *t_40_* = −7.44, *p*<0.0001). This suggests target streaming was more challenging under conditions of linguistic (i.e., speech-on-speech) compared to energetic (i.e., speech-on-noise) masking loads.

For speed, we found main effects of masker count [*F_1,140_*=33.46, *p*<0.0001, 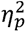*=* 0.42] and masker direction [*F_1,140_* =26.33, *p*<0.0001, 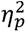*=* 0.16] on speech recognition RTs (**Fig. 3b**). These data reveal that decision speeds were predictably slower in more challenging multi-talker scenarios and with an increasing number of competing talkers.

Localization errors are shown in **Figure 3c**. Listeners localized targets within ∼2-3 speakers (40-60° error). Localization varied with both masker count and direction [interaction: *F_3,140_* =12.28, *p*<0.0001, 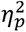*=* 0.21]. Tukey contrasts show the interaction was attributable to masker-related differences at 2 and 4 maskers counts. This suggests the influence of masker content (i.e., whether competing talkers were intelligible or not) was prominent only at higher talker counts.

Source monitoring is shown in **Figure 3d**. In general, listeners could distinguish how many talkers were in the soundscape with up to ∼3 simultaneous voices. Performance plateaued thereafter suggesting a saturating effect in source monitoring performance. This was confirmed by a sole main effect of masker count [*F_3,140_* =606.41, *p*<0.0001, 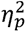*=* 0.93]. The lack of masker direction effect indicates source monitoring did not depend on masker intelligibility.

### 3.3 Phoneme categorization

Phoneme categorization for CVs and vowels under the 2AFC vs. VAS task is shown in **Figure 4**. Identification slopes, reflecting the degree of categoricity in listener response pattern, were modulated by an interaction between stimulus continuum and task [*F_1,63_* =4.47, *p*=0.038, 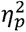*=* 0.07]. Multiple comparisons revealed this interaction was due to steeper identification for vowels compared to CVs but only in the 2AFC task (**Fig. 4a**). Slopes were invariant under VAS labeling (**Fig. 4c**). However, the stimulus effect is not evident under tasks which promote continuous/gradient modes of listening, as in the VAS paradigm.

RT labeling speeds are shown in **Fig. 4b** and **4d**. RTs were ∼750 ms later when categorizing speech sounds under VAS compared to 2AFC labeling [*F_1,394.3_*=1090.4, *p*<0.0001, 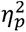 *=* 0.73]. However, this effect is largely expected due to trivial differences in the nature of the motor response in the 2AFC vs. VAS tasks (i.e., keyboard vs. mouse). Consequently, we normalized RTs by subtracting the mean across tokens to highlight the *relative* changes in speed between continua and tokens (Bidelman et al., 2019). An ANOVA conducted on RTs revealed main effects of token [*F_4,394.3_*=2.48, *p=*0.043, 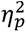 *=* 0.02] and stimulus [*F_1,394.3_* =12.83, *p=*0.0004, 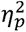 *=* 0.03]. The stimulus effect was due to slightly faster (∼70 ms) RTs for vowels compared to CVs. The token effect was attributable to the hallmark slowing (i.e., inverted-V pattern) in labeling speeds near the ambiguous midpoint of the continuum for vowels in both tasks [2AFC: *t_414_* = 2.56, *p* = 0.011; VAS: *t_414_* = 2.36, *p* = 0.0187] (Pisoni and Tash, 1974; Bidelman and Walker, 2017; Carter and Bidelman, 2021). However, this slowing effect due to phonetic ambiguity was not observed for CVs under either task (*p*s > 0.29), consistent with prior work (Carter and Bidelman, 2021; Carter et al., 2022). These data support the notion that CVs are heard more categorically and with lesser phonetic ambiguity than vowels (Pisoni, 1973; Altmann et al., 2014; Carter et al., 2022). They also suggest the nature of the task changes categorization outcomes, with a 2AFC task structure producing more categorical/discrete hearing than a VAS task structure.

### 3.4 Relations between listening categorization and cocktail party SIN perception

Our phoneme labeling tasks were designed to promote more discrete (2AFC) vs. gradient (VAS) hearing. In particular, VAS ratings are thought to better isolate continuous vs. categorical modes of speech perception at the individual level (Kapnoula et al., 2017). To quantify such individual differences in listening strategy, we divided our sample into “discrete” vs. “continuous” categorizers based on the distribution of their VAS labeling and Hartigan’s Dip statistic (Hartigan and Hartigan, 1985). The Dip metric tests the intensity of bimodality of the data and thus whether labeling reports are bimodal (high dip score =categorical) or unimodal (low dip score=continuous) (**Fig. 5**).

**Figure 5:**
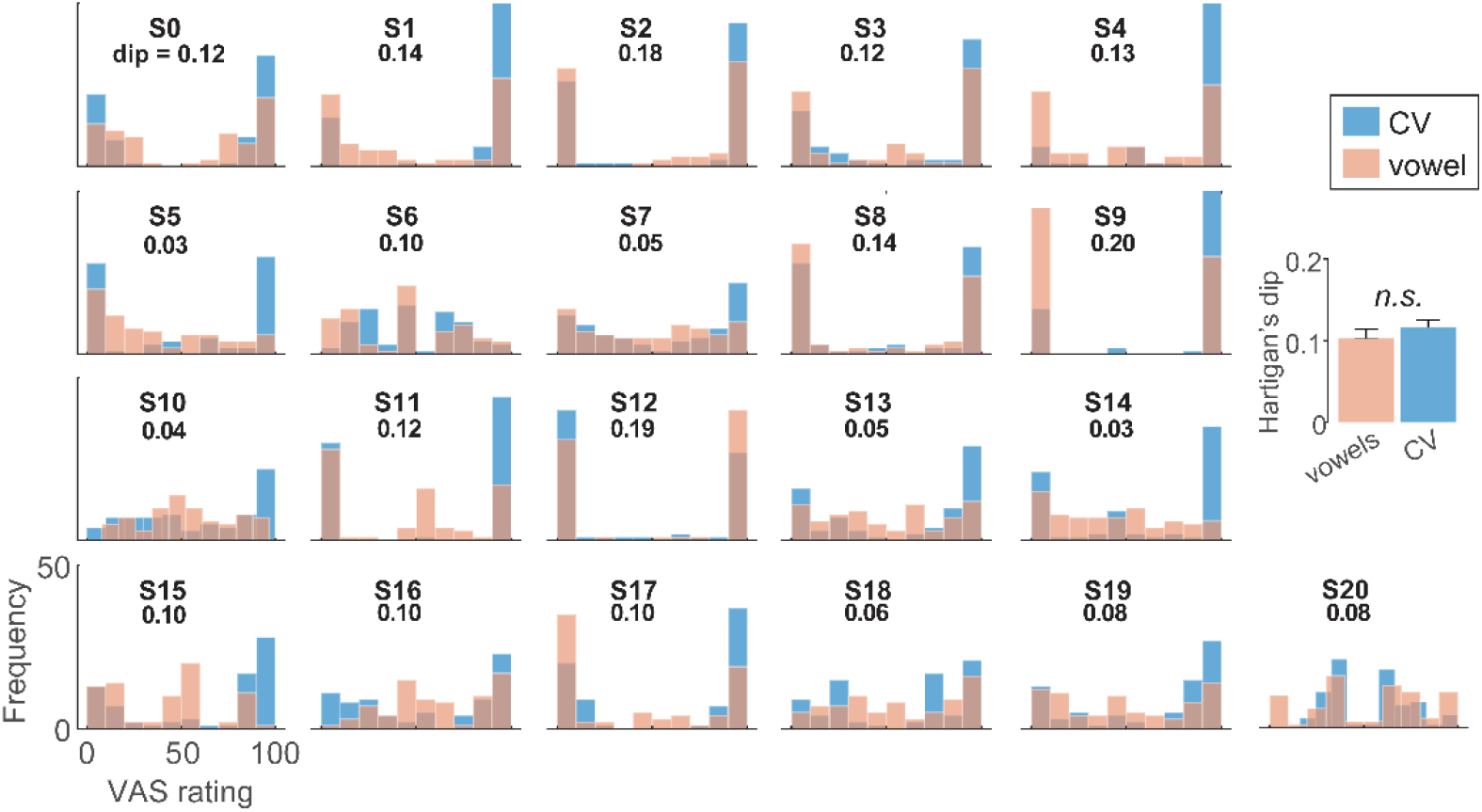
VAS ratings reveal stark individual differences in categorization and “continuous” vs. “categorial” listeners. Individual histograms show the distribution of each listeners phonetic labeling for CV and vowel sounds. Discrete (categorical) listeners produce more binary categorization where responses lump near endpoint tokens (e.g., S2). In contrast, continuous (gradient) listeners tend to hear the continuum in a gradient fashion (e.g., S16). Inset values show Hartigan’s Dip statistic (Hartigan and Hartigan, 1985) score, quantifying the bimodality—and thus categoricity—of each distribution. Higher dip values= discrete categorization; low values = continuous categorization. (inset) Dip values are similar between CV and vowels suggesting it is a reliable measure of listener strategy that is independent of speech material. errorbars = ± 1 s.e.m.

Being a discrete/continuous categorizer did not depend on speech content as Hartigan’s Dip statistic was similar between CVs and vowels [*t_20_* = −1.15, *p* = 0.26]. This suggests it was a reliable profile of individual listener strategy that is independent of speech material. Given there were no stimulus-related differences in dip scores, we pooled CV and vowel VAS data for subsequent analyses. We then divided the sample into two groups based on whether an individual’s dip statistic computed from their VAS ratings showed significant (*p*<0.01) evidence of bimodality. This resulted in two groups: “discrete” (n=14) vs. “continuous” (n=7) listeners.

**Fig. 6a** shows cocktail party speech recognition performance (as in Fig. 3a) split by group. For each listener, we computed the degree of informational masking (IM) experienced in the speech streaming task, measured as the difference in recognition performance (raw %-correct scores) in the forward and reverse masker conditions at each masker count (**Fig. 6a**). The rationale behind this metric is that speech-on-speech masking in the forward talker condition contains additional linguistic interference due to the intelligibility of the masking talkers that further hinders figure-ground speech perception (Swaminathan et al., 2015; Yoo and Bidelman, 2019).

**Figure 6:**
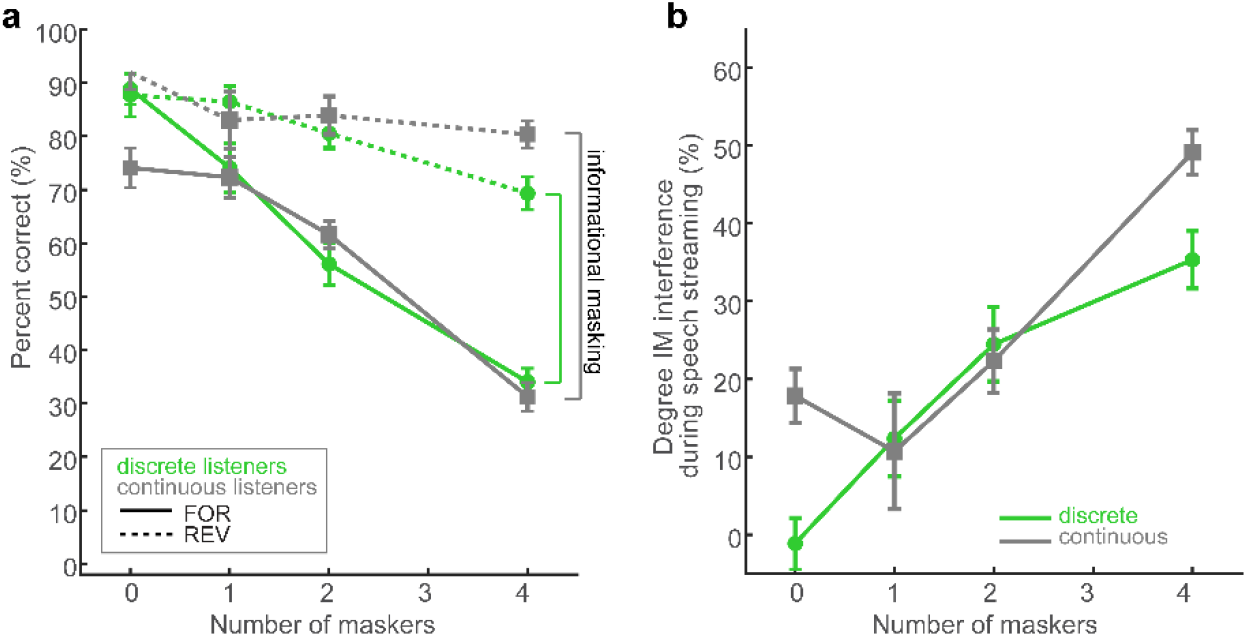
Categorical listeners are less susceptible to speech interference at the “cocktail party.” **(a)** Speech recognition performance in the streaming task for discrete and continuous listeners. Otherwise as in Fig. 3. Listener strategy was determined via Hartigan’s dip statistic (Hartigan and Hartigan, 1985) applied to VAS labeling (i.e., Fig. 5) to identify individuals with bimodal (categorical) vs. unimodal (continuous) response distributions. Information masking (IM) in speech streaming was measured as the difference in recognition performance between forward and reverse masker conditions at each masker count. (**b**) Categorical listeners show less IM during speech streaming than their continuous listener peers. errorbars = ±1 s.e.m.

**Fig. 6b** shows IM computed for “discrete” vs. “continuous” listeners. A 2-way ANOVA conducted on IM revealed main effects of masker count [*F_3,76_* =19.89, *p*<0.0001, 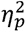 *=* 0.44] and group [*F_1,76_* =4.43, *p*=0.038, 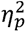 *=* 0.06] with no interaction [*F_3,76_* =2.45, *p*=0.07, 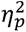 *=* 0.09] (**Fig. 6b**). The masker effect was due to a steady and expected increase in IM with increasing masker talker counts. More critically, the main effect of group indicates categorical listeners were less susceptible to IM during cocktail party speech perception than their gradient peers across the board.

### 3.5 Relations between EHFs and SIN

Correlations between QuickSIN and speech streaming measures were insignificant (all *p*s > 0.24), suggesting they tap different factors of auditory figure-ground processing. Similarly, QuickSIN was not related to any of the phoneme categorization measures (all *p*s > 0.14).

Despite all listeners having normal hearing, EHF thresholds did predict QuickSIN performance (Pearson’s *r* =0.48, *p*=0.0259). Slightly worse (though still within normal limits) high-frequency hearing sensitivity was associated with poorer (i.e., larger) QuickSIN scores. However, EHFs were not related to any measures of streaming performance (all *p*s > 0.05), indicating cocktail party perception was independent of high-frequency hearing.

## 4. Discussion

By measuring phoneme identification and degraded speech recognition in a multi-talker soundscape, we establish a new link between two fundamental operations in speech processing: categorization and speech-in-noise (SIN) perception. Our findings suggest the mere process of binning speech sounds into categories provides a robust mechanism to aid perception at the “cocktail party” by fortifying abstract categories from the acoustic signal and making the speech code more resistant to external interferences.

### 4.1 Speech recognition at the cocktail party: accuracy, speed, localization, and source monitoring

Our cocktail party speech task revealed that the ability to stream target speech amidst concurrent talkers depends critically on the linguistic nature of the maskers (i.e., whether or not they are interpreted as speech). Recognition accuracy and speed expectedly declined with increasing multi-talker interferers (Bidelman and Yoo, 2020). Poorer speech recognition with additional talkers is consistent with a reduction in spatial release from masking as more concurrent streams reduce the separability of the target in the soundfield (Pastore and Yost, 2017). More limited performance at higher masker counts also agrees with previous behavioral studies which show spatial release from masking is effectively limited to fewer than 6 sound sources (Yost, 2017).

Performance was also better overall during reversed compared to forward maskers. This effect was also anticipated and can be explained by the fact that forward masks contain additional informational masking (IM) due to the linguistic information of speech-on-speech masking. In contrast, reverse masks are not interpreted as intelligible speech, *per se*, and contain only energetic masking (EM). While EM is related to the interference of cochlear excitation patterns of the signal and masker and thus, peripheral hearing function, IM reflects additional confusability of the signals and thus represents central-cognitive aspects of figure-ground perception (Moore, 2012). Consequently, the forward talker condition containing speech-on-speech masking is more difficult given the added challenge of parsing multiple linguistic signals (Swaminathan et al., 2015; Yoo and Bidelman, 2019).

In terms of localizing and monitoring talkers in the acoustic environment, we found listeners pinpointed targets within ∼2-3 speakers (40-60° error), consistent with our previous auditory streaming studies (Bidelman and Yoo, 2020). However, localization showed an interaction effect, suggesting the influence of masker content (i.e., whether competing talkers were intelligible or not) was more prominent only at higher talker counts. One explanation for this effect is that the localization task was delayed compared to recognition. There is evidence listeners can localize sound sources even if they cannot identify them (Rakerd et al., 1999). Indeed, determining *where* a signal is emitted in the soundscape has a clear biological advantage over identifying *what* it is. Relatedly, our source monitoring results demonstrate that listeners are only able to identify the presence of ∼3 talkers in the soundscape, despite more being present in the environment. This indicates a capacity limit in auditory streaming whereby listeners can only resolve up to ∼3 distinct voices at any one time (present study; Yost et al., 2019). This finding is also consistent with channel capacity limits in auditory processing and notions that listeners cluster task-irrelevant sounds (e.g., background talkers) into a single stream to improve the perceptual segregation and identification of target information (Alain and Woods, 1994; Yost et al., 2021).

### 4.2 Speech recognition in noise partially relates to EHF thresholds

Prior studies have suggested SIN perception is related to high-frequency hearing sensitivity, as measured via EHF thresholds, even in “normal hearing” individuals (Monson et al., 2019; Mishra et al., 2022). In the present study, we similarly observe a link between EHF audiometric thresholds and QuickSIN scores. Slightly worse (though still within normal limits) high-frequency hearing sensitivity was associated with poorer (i.e., larger) QuickSIN scores. Though we note EHF thresholds did not predict cocktail party streaming. The link between some SIN measures and EHFs is consistent with some (Monson et al., 2019; Mishra et al., 2022) though not all studies (cf. Liberman et al., 2016; Lai and Bidelman, 2022). Additional work is needed to understand putative relationships between high-frequency hearing and SIN abilities (even in normal hearing ears).

### 4.3 Categorization skills are related to SIN processing

VAS ratings of speech-sound continua allowed us to isolate continuous vs. categorical modes of speech perception and quantify individual differences in listening strategy based phoneme labeling skills (Kapnoula et al., 2017). Applying this approach, we show listeners can be reliably pooled into “discrete” vs. “continuous” categorizers based on the distribution of their phoneme labeling. This division was not idiosyncratic to the specific speech content (i.e., whether listeners are identifying CVs or vowels), suggesting the behavioral profiles are a reliable index of individual listener strategy. Relevant to our hypothesis of a categorization-SIN relation was listeners’ performance on the cocktail party and QuickSIN tasks as a split of these functional differences in perceptual identification strategy.

Measuring the degree of informational masking experienced by listeners in speech streaming, we found SIN recognition was robustly predicted by categoricity in hearing. While IM expectedly increased for all listeners with increasing talker count, interestingly, “discrete” listeners showed less speech-on-speech hinderance in performance than their “continuous” hearing peers. This group effect indicates that certain listeners who hear speech sounds in a more categorical manner are less susceptible to interference at the cocktail party. That a discrete listening strategy is more beneficial to complex SIN processing is at odds with prior work implying a benefit of gradient listening (McMurray et al., 2008; Kapnoula et al., 2017). However, when put to empirical scrutiny, studies have failed to establish a consistent pattern between SIN performance and listening strategy. For example, word comprehension in noise for garden path and AzBio sentences do not correlate with listening strategy measured by VAS categorization (Kapnoula et al., 2017; Kapnoula et al., 2021). These findings, coupled with current results, suggest that while listeners can maintain access to continuous, within-category cues (Pisoni and Lazarus, 1974; Pisoni and Tash, 1974; Spivey et al., 2005; Huette and McMurray, 2010; Bidelman and Carter, 2023), it is not generally beneficial to parsing noise-degraded speech. Instead, our data support the notion that hearing speech in a categorical mode (a more abstract level of coding; Pisoni and Lazarus, 1974) aids degraded speech perception (e.g., Bidelman et al., 2019; Bidelman et al., 2020; Lewis and Bidelman, 2020; Bidelman and Carter, 2023). Presumably, categoricity allows for irrelevant variations in the physical surface features of the signal to be largely discarded in favor of retaining the abstract and robust phonetic code for speech that is more impervious to noise interference (Bidelman et al., 2020).

Our results corroborate notions that category-level cues provide easier readout to brain processing (Pisoni and Tash, 1974; Guenther et al., 2004; Bidelman et al., 2013; Reetzke et al., 2018; Bidelman et al., 2020) and notions that categorical percepts are more impervious to surface-level degradations that can corrupt speech recognition (Gifford et al., 2014; Helie, 2017; Bidelman et al., 2019; Bidelman et al., 2020). Previous studies comparing phoneme categorization performed under clean vs. noise-degraded listening conditions reveals listeners easily label speech even at unfavorable SNRs (Bidelman et al., 2019; Bidelman et al., 2020). Categories might also aid the extraction of target speech percepts from interfering sound sources by reducing listening effort. This notion is supported by behavioral and physiological data (ERP: Bidelman et al., 2020; pupillometry: Lewis and Bidelman, 2020). Relatedly, perceptual warping effects in speech categorization (Ganong, 1980; Tuller et al., 1994; Myers and Blumstein, 2008; Bidelman et al., 2021; Carter et al., 2022)—where tokens can be made to sound closer to distal prototypes in acoustic-phonetic space—are more prominent under noise relative to clean speech (Bidelman and Carter, 2023). Indeed, in mousetracking studies on phonetic categorization, listeners take a more direct and faster motor path when classifying sounds amidst noise (Bidelman and Carter, 2023). This could result from stronger perceptual attraction to category members (Carter et al., 2021) or reduced decision ambiguity (Viswanathan and Kelty-Stephen, 2018) supplied by the reductionist process of category mapping.

### 4.4 Categorization is related to discreteness/gradiency rather than noisy perception

Categorization is typically quantified by the slope of listeners’ identification functions in a 2AFC task. However, shallower slopes in a 2AFC task may reflect perceptual gradiency and/or more internal noise in cue encoding. Both factors would tend to flatten a sigmoidal identification curve and thus are conflated in binary 2AFC tasks. Consequently, it has been argued that VAS labeling provides a purer measure of categorization discreteness/gradiency that is immune to the effects of sensory noise in behavior (Kapnoula et al., 2017; Apfelbaum et al., 2022). The confounding of categoricity and sensory noise was also our primary motivation for using the Dip statistic (Hartigan and Hartigan, 1985) to define “categorical” vs. “continuous” listeners rather than identification slopes.

Still, to test the hypothesis that psychometric slopes reflect perceptual categoricity rather than internal decision noise, we estimated the noise in the VAS responses, measured as the *SD* in labeling reports across tokens (e.g., Kapnoula et al., 2017). Pooling across CV and vowel data, we found 2AFC slopes were not correlated with noise in the VAS task [*r =* 0.06, *p* = 0.79]. These findings thus do not support the assertion that shallower slopes (i.e., weaker categorization) in a 2AFC task is due to increased internal sensory noise (cf. Kapnoula et al., 2017). More critically, we found no correspondence between Dip statistic scores (bimodality of responses) and response noise [*r* = −0.06, *p* = 0.79]. Thus, our data suggest the slopes in 2AFC and VAS categorization tasks reflect the degree to which sounds are heard categorically rather than noisier responding, *per se*.

### 4.6 Broader implications of a categorization-SIN link

SIN performance has long been linked to higher level cognitive skills—most notably, working memory (Tamati et al., 2013; Füllgrabe and Rosen, 2016; Kapnoula et al., 2017; Bidelman and Yoo, 2020). Our findings establish a new link between two fundamental and arguably more rudimentary *perceptual* operations (categorization, figure-ground) that could explain broader individual differences in SIN skills among normal and clinical populations alike. For instance, the degree to which listeners show categorical vs. gradient perception might reflect the strength of phonological processing, which could have ramifications for understanding both theoretical accounts of speech perception and certain clinical disorders that impair sound-to-meaning mapping (e.g., dyslexia; Werker and Tees, 1987; Joanisse et al., 2000; Calcus et al., 2016). It has even been suggested that deficits in speech categorization among certain developmental disorders might also be more prominent in noise (Calcus et al., 2016). Both categorization and speech-in-noise aspects of hearing show considerable *inter-*subject (but less *intra*-subject) variability (present study; Song et al., 2011; Billings et al., 2013; Bidelman et al., 2018; Bidelman and Momtaz, 2021; Carter et al., 2022). Thus, it is tempting to infer that figure-ground deficits observed in some auditory and language-based learning disorders (Cunningham et al., 2001; Warrier et al., 2004; Putter-Katz et al., 2008; Lagacé et al., 2010; Dole et al., 2012; Dole et al., 2014) result from a failure to flexibly warp category representations of the speech code. On one hand, graded/continuous perception might be advantageous for speech perception in noise since it would allow listeners access to all acoustic information in the signal, potentially allowing them to “hedge” their bets on what they are hearing in face of ambiguity (Kapnoula et al., 2017). On the other hand, if a large portion of the perceptual space is corrupted by noise, hearing in discrete units might be preferrable to allow category members to “pop out” among the noise and facilitate speech processing (Nothdurft, 1991; Pérez-Gay Juárez et al., 2019; Bidelman et al., 2020). Our data here lead us to infer that the discretization of auditory information is more beneficial to parsing speech in realistic cocktail party SIN scenarios and how well a listener can extract (or suppress) concurrent speech information. Nevertheless, future studies in clinical populations are needed to determine if SIN deficits commonly observed in clinical disorders truly result from deficits in sound-to-label mapping (i.e., categorization).

## Acknowledgements

The authors thank Rose Rizzi, Jessica MacLean, Jack Stirn, Elizabeth Drobny, and Serenity Seigel for comments on earlier versions of the manuscript. Requests for data and materials should be directed to G.M.B.

## Footnotes

1 The University of Memphis anechoic chamber is a room-within-a room design featuring a 24’ x 24’ x 24’ IAC chamber with floor/wall/ceiling Metadyne® acoustic wedge coverage. The noise lock provides an STC 61 noise rating (low cutoff frequency=100 Hz). A 36 channel Renkus-Heinz speaker array surrounds the seating location (16 were used in the experiment). Multichannel audio control is achieved by a TDT RX8 Multi-I/O Processor (Tucker Davis Technologies). Six Focusrite and Ashley Ne8250 amplifiers drive the speakers via a RedNet Dante MADI interface.

